# VaxSeer: Selecting influenza vaccines through evolutionary and antigenicity models

**DOI:** 10.1101/2023.11.14.567037

**Authors:** Wenxian Shi, Jeremy Wohlwend, Menghua Wu, Regina Barzilay

## Abstract

Current vaccines provide limited protection against rapidly evolving viruses. For example, the flu vaccine’s effectiveness has averaged below 40% for the past five years. Today, clinical outcomes of vaccine effectiveness can only be assessed retrospectively. Since vaccine strains are selected at least six months ahead of flu season, prospective estimation of their effectiveness is crucial but remains under-explored. In this paper, we propose an *in-silico* method named VaxSeer that selects vaccine strains based on their coverage scores, which quantifies *expected* vaccine effectiveness in future seasons. This score considers both the future dominance of circulating viruses and antigenic profiles of vaccine candidates. Based on historical WHO data, our approach consistently selects superior strains than the annual recommendations. Finally, the prospective coverage score exhibits a strong correlation with retrospective vaccine effectiveness and reduced disease burden, highlighting the promise of this framework in driving the vaccine selection process.

## 1 Main

The genetic composition of an influenza vaccine is one of the main drivers of its effectiveness. Twice a year, a panel of experts from the World Health Organization (WHO) gathers to recommend vaccine strains for the upcoming flu seasons. Their goal is to identify vaccine strains that will induce antibodies effective against viruses expected to circulate in the upcoming seasons. If their recommendations are well-aligned with the future viral landscape, the effectiveness^1^ of an inactivated influenza vaccine may reach up to 40% to 60% in a given season [2]. However, despite decades of research in prevention and surveillance, current influenza vaccines provide limited protection: the Centers for Disease Control and Prevention (CDC) report that vaccine effectiveness in the United States has dropped below 40% in five of the past ten years [3–7]. In fact, during the 2014-2015 winter season, vaccine effectiveness was only 19% [3].

Currently, vaccine effectiveness and related clinical outcomes, such as reduction of hospitalization and mortality, are measured retrospectively, after the conclusion of each flu season [1]. During the vaccine design process, however, it is critical to develop a proxy for each vaccine’s *expected* effectiveness in the upcoming season. There are two primary barriers to accurate vaccine strain selection: the experimental burden of testing vaccine and virus combinations, and the inherent uncertainty of future viral mutations. First, it is prohibitively expensive to validate all candidate vaccines against all circulating viruses, and there are limited viral specimens available for study. *In-vitro* assays such as the hemagglutination inhibition (HI) test [8] are commonly used to assess the extent to which a vaccine’s induced antibodies inhibit a specific virus. However, these experiments can only be run on a limited number of vaccine candidates, typically fewer than ten [9]. Second, inactivated influenza vaccines require a production period of six to nine months [10], during which the distribution of virus strains may change significantly [11–13] (illustrated in Fig. S2). As a result, strains prevalent during the actual flu season may exhibit different antigenic profiles than expected, reducing the effectiveness of antibodies elicited by the selected vaccines. Therefore, our hypothesis is that simultaneously modeling the antigenic similarity of vaccine strains against virus strains and the future distribution of viruses can lead to more accurate estimates of a vaccine’s expected effectiveness.

This paper presents a computational approach named VaxSeer that realizes this idea in the context of influenza. We aim to maximize the *coverage score*, a surrogate metric that prospectively estimates vaccine effectiveness by considering the *antigeniticy* of a vaccine to a virus strain and the *dominance* of future virus strains. To evaluate the antigenic similarity between vaccines and circulating strains, we define antigenicity as the inhibition capacity of antibodies produced by a vaccine to a specific virus. The holistic antigenicity of the vaccine strain concerns multiple circulating viruses, in the context of their future dominance. Dominance is defined as the frequency of occurrence during a particular season. In the case of influenza, for instance, we can quantify antigenicity through the outcomes of HI tests [8]. We separately train two models to predict each of these respective values. When combined, these models allow us to compute the predicted coverage score, a vaccine’s expected effectiveness over a panel of potential circulating viruses. Compared to existing assays or metrics, which model individual aspects of immunology like viral fitness [14–20] and antigenic properties [21–24], our goal is to optimize a holistic score that prospectively quantifies the effectiveness of vaccines. Retrospective analyses show that the coverage score is strongly correlated with vaccine effectiveness and reduction of disease burden. Thus, selecting vaccine strains based on coverage score has the potential to improve clinical endpoints of vaccine effectiveness. Finally, this approach can also be applied to any other circulating pathogen with sufficient antigenic and evolutionary data, such as SARS-CoV-2.

## Results

### Overview of VaxSeer

The coverage score is a prospective surrogate for vaccine effectiveness that quantifies the antigenicity of a vaccine against viral strains, weighted by their future dominance. Specifically, let 𝒳 denote the set of circulating viruses, and let *v* denote the vaccine in question. We define the coverage score *CS* of a vaccine *v* during a future season *t* as

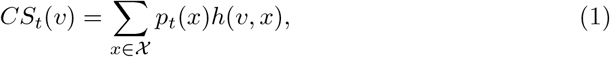

where *p*_*t*_(*x*) is the dominance of virus *x* in season *t*, and *h*(*v, x*) measures the antigenicity between vaccine *v* and virus *x* (Fig. 1a). Large values of *p*_*t*_(*x*) indicate that virus *x* has high dominance, and large values of *h*(*v, x*) indicate that vaccine *v* is effective against virus *x*. We show that coverage score is positively correlated with vaccine effectiveness (Fig 1d). The vaccine candidates with the highest (predicted) coverage scores are those recommended by our algorithm.

**Fig. 1:**
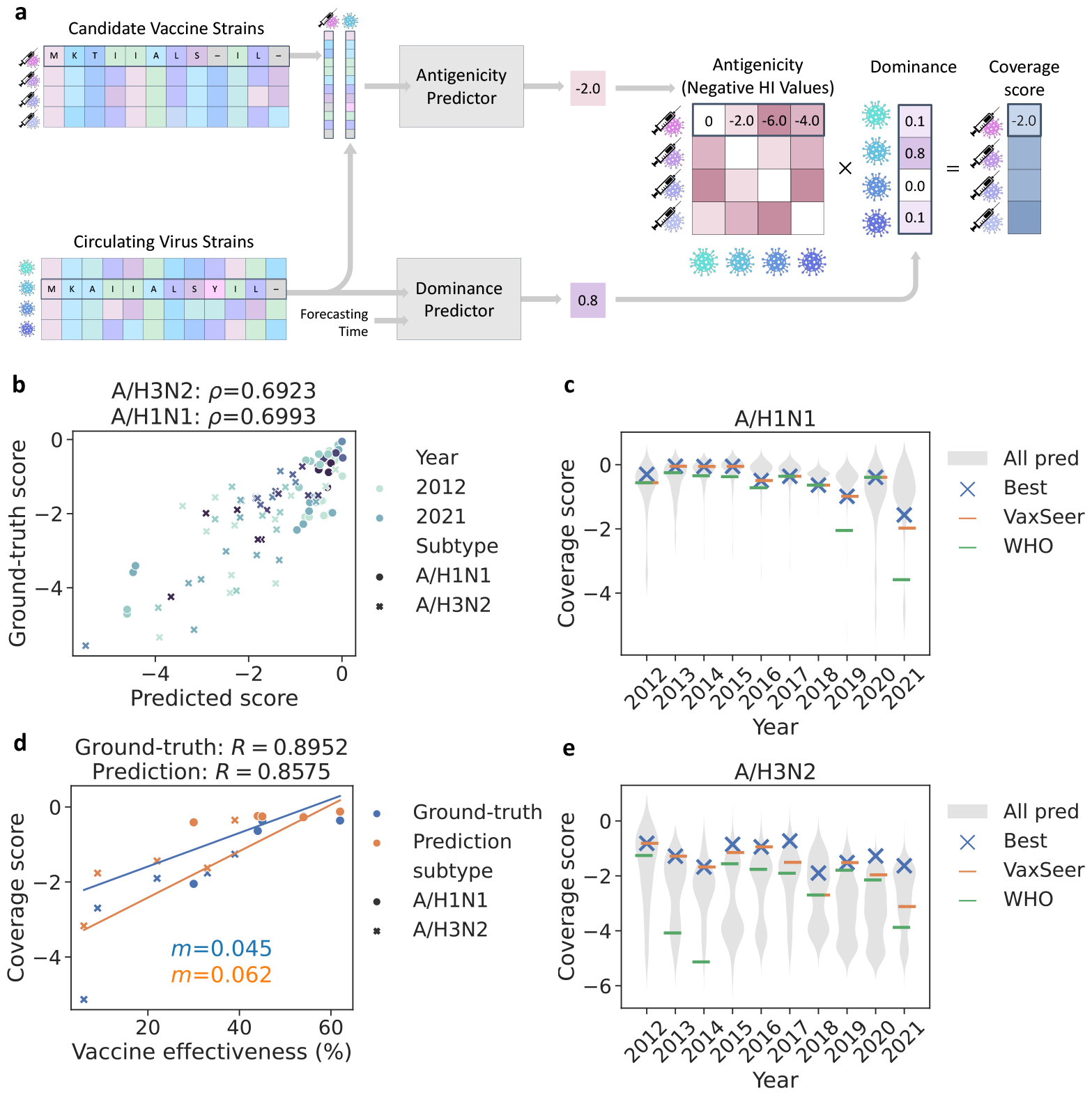
Overview of VaxSeer for vaccine selection. (**a**) A two-track model for predicting the coverage score. We combine the dominance and antigenicity predictor’s outputs to estimate the coverage score. (**b**) Predicted coverage score demonstrates statistically-significant correlation with ground-truth coverage score (Spearman rank correlation *ρ* with P-value *P <* 0.005). (**c, e**) Vaccine strains selected by VaxSeer have higher ground-truth coverage scores than those recommended by the WHO. VaxSeer frequently selects the strains with the *highest* experimentally-measured coverage scores. Violin plots depict the distribution of predicted coverage scores among all candidate vaccines, including those with low experimental coverage. (**d**) Both ground-truth and predicted coverage scores exhibit strong correlations with the real-world vaccine effectiveness, estimated by the CDC (Pearson correlation *R* with P-value *P <* 0.005, linear regression slope *m*).

Due to the difficulty of forecasting future viral dominance and the costs associated with large-scale antigenicity assays, we propose to predict the coverage score with an *in-silico* approach (Fig. 1a). In our influenza case study, we represent vaccine and virus strains by their hemagglutinin (HA) protein sequences, which contain the major antigens targeted by antibodies [25, 26]. The antigenicity *h* is defined based on the HI test assay [8]. In particular, we set the *h* equal to the negative HI value, given by the HI titer between vaccine *v* and virus *x* [8]. Given the HA protein sequences of a vaccine and a virus, the *antigenicity predictor* regresses their HI test result (antigenicity) by aligning the two sequences and modeling the antigenic similarity between residues across sequences. Given the HA protein sequence of a virus, the *dominance predictor* outputs a dynamic, time-resolved probability that it will be isolated in a future flu season (dominance). These learning modules can capture epistatic relationships.

We conduct retrospective validation of our model on flu surveillance data and antigenic analysis data from 2012 to 2021. It is possible to compute the *ground-truth* coverage score from experimental HI test results and (retrospective) flu surveillance data. Fig. 1b shows that our model’s predicted coverage scores are highly consistent with the ground-truth coverage scores. Furthermore, our predicted coverage score enables the selection of better vaccine strains than the WHO’s recommendations, as measured by ground-truth coverage score (Fig.1c, Fig. 1e). We also show that both the predicted and ground-truth coverage scores of each year’s recommended vaccines correlate strongly with their measured effectiveness (Fig. 1d). The linear fitting suggests that a 0.62 increase in predicted coverage score corresponds to a 10% increase in vaccine effectiveness. Finally, coverage score is also strongly correlated with clinical endpoints that quantify the reduction of disease burden (Fig. 3c-3f). Therefore, optimizing vaccines to maximize coverage score not only enables us to improve vaccine effectiveness, but also reduces the potential disease burden of influenza.

### Experimental setup

We conducted retrospective validation of VaxSeer on the A/H1N1 and A/H3N2 sub-types of influenza. To quantify viral dominance during each season, we downloaded influenza HA protein sequences with associated collection times from GISAID [27]. For antigenicity measurements, we collected HI test results from WHO reports^2^ prepared for the annual consultations on the composition of influenza vaccines from 2003 to February 2023. Finally, influenza vaccine effectiveness and clinical endpoints were gathered from annual CDC publications [3–7].

Our evaluation spans the 2012 to 2021 winter seasons (October to March). In line with the WHO’s recommendation schedule, we train our models on strains collected and HI tests conducted up to eight months prior to the start of the season (the preceding February). To construct the sets of candidate vaccine strains and circulating virus strains, we consider all virus strains that were isolated at least five times within the past three years. For example, to evaluate the 2021-2022 winter season (October 2021 to March 2022), we train our models on any data collected before February 2021, and we compute coverage scores over strains isolated between February 2018 and February 2021.

Ground-truth scores are computed retrospectively, using data collected during the *actual* flu season (October to March). Each flu strain’s ground-truth dominance is equal to its frequency of occurrence (the number of times each strain was isolated, divided by the total number of strains isolated during the season). The ground-truth coverage score of a vaccine is determined by taking the dominance-weighted average of antigenicity over all viruses. To ensure sufficient coverage of circulating viruses, we only computed ground-truth coverage scores for vaccines with HI test results against at least 40% of circulating viral sequences during the season of interest. A total of 51 candidate vaccines for A/H3N2 and 50 candidate vaccines for A/H1N1 were subjected to comparison.

### Vaccine selection based on coverage score

As Fig. 1b shows, there is a strong correlation between our predicted scores and the ground-truth scores for all tested vaccines. Based on ground-truth coverage score, VaxSeer selects better vaccine strains than the WHO’s past recommendations in six out of ten years for A/H1N1 and nine out of the ten years for A/H3N2 (Fig. 1c, 1e). In fact, VaxSeer successfully recommends the *best* vaccine strain (top ground-truth coverage score) in eight out of the ten years for A/H1N1 and five out of ten years for A/H3N2, whereas the WHO’s recommendation matches the best vaccine strain in just three out of ten for A/H1N1 and zero out of ten for A/H3N2. Due to WHO experimental constraints, only a subset of candidate vaccines are tested broadly (over 40% of influenza viruses). Thus, we can only compute ground-truth coverage scores over this subset, and the strains selected in Fig. 1c and Fig. 1e only reflect these limited options. If we look at the distributions of predicted coverage scores over *all* candidate vaccine strains (violin plots in Fig. 1c and 1e), there are strains that scored even higher, but were not subject to sufficient experimental validation by WHO. This highlights the possibility that there may exist even more efficacious vaccine strains waiting to be discovered.

Fig. 2a-2b and Fig. 2d-2e present four specific case studies: the 2016 and 2019 seasons for A/H3N2 and A/H1N1. Each point represents a candidate vaccine. The green dots are the strains recommended by the WHO, while the orange dots are those recommended by VaxSeer. These figures illustrate a strong correlation between the scores predicted by our model (x-axis) and the ground-truth scores (y-axis). In particular, in Fig. 2d, though the WHO recommendation (A/California/07/2009) has a decent ground-truth coverage score, VaxSeer proposes an alternative with an even higher coverage score (A/Michigan/45/2015, collected earliest on 2015-09-07). Promisingly, A/Michigan/45/2015 was recommended by the WHO for the subsequent winter season.

**Fig. 2:**
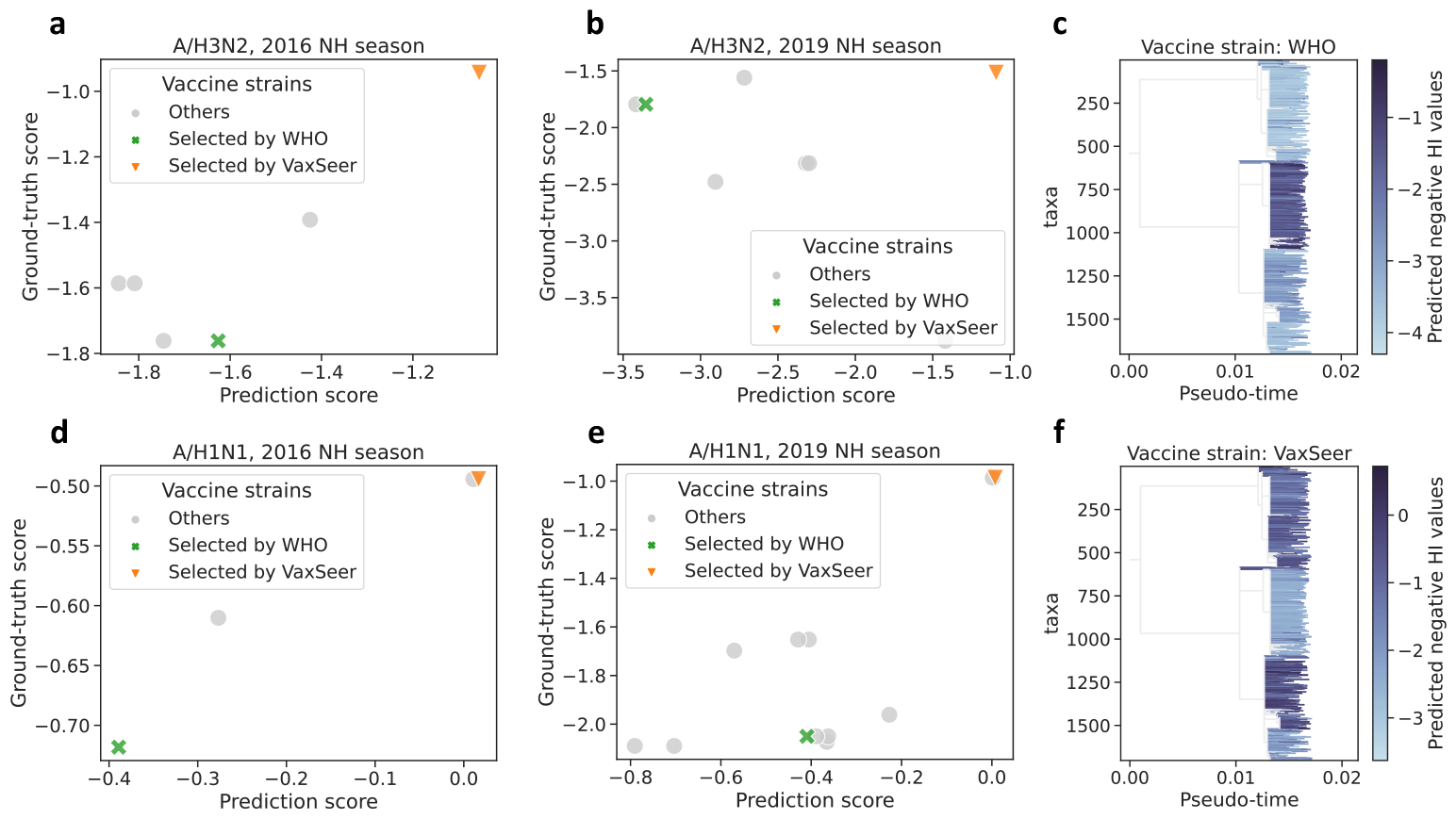
(**a, b, d, e**) Ground-truth coverage scores vs. predicted coverage scores, high-lighting WHO recommendations (green) and VaxSeer’s top-ranked candidate (orange). (**c, f**) The predicted antigenicity (negative HI values) of circulating A/H3N2 strains in the 2019 NH season, with respect to the vaccine (**c**) recommended by WHO and (**f**) VaxSeer. Our recommended vaccine covers a larger variety of circulating viruses than the WHO’s recommendation.

**Fig. 3:**
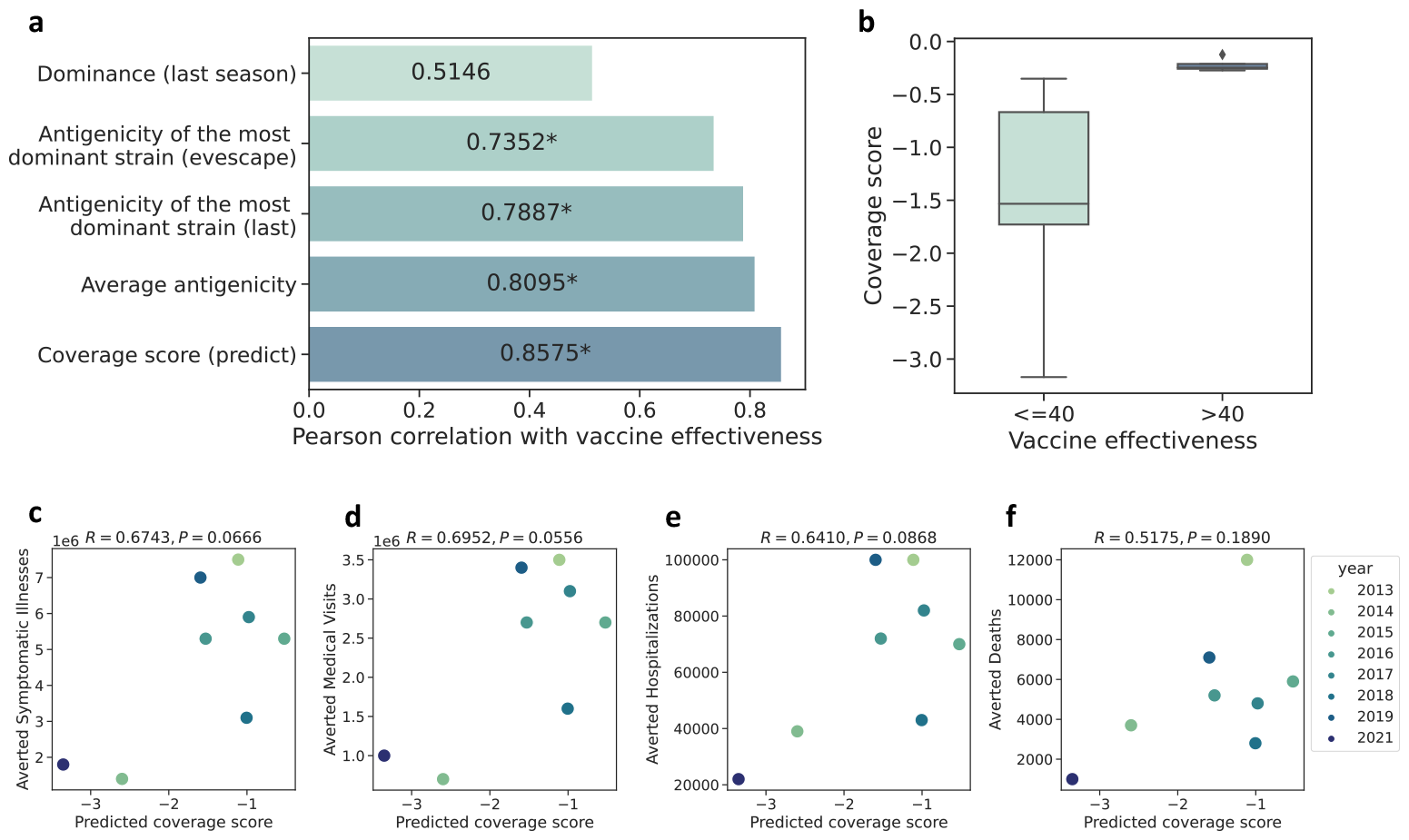
The predicted coverage score is correlated with vaccine effectiveness and disease burden prevention. (**a**) The Pearson correlation between vaccine effectiveness and various scoring strategies for the vaccine strains. The star mark (*) indicates *P <* 0.05. Our coverage score shows the strongest correlation. (**b**) The vaccines with higher effectiveness (*>* 40) have higher coverage scores than vaccines with lower effectiveness (*≤* 40). The medians of the coverage scores are depicted by the center lines within the boxes. The box spans from the first to the third quartile, and the whiskers cover 1.5 times the interquartile range. Outliers beyond the whiskers are represented by individual marks. (**c-f**) Correlations between the predicted coverage scores and the estimated number of influenza illnesses, medical visits, hospitalizations, and deaths averted by vaccination. Pearson correlations (*R*) and corresponding P-values (*P*) are illustrated in the text.

To visualize the breadth in coverage of different vaccine strains, we plot the predicted antigenicity (negative HI values) of A/H3N2 strains circulating in 2019 winter season. The strain recommended by the WHO only aligns with one circulating clade (Fig. 2c). In contrast, the strain recommended by VaxSeer offers a complementary antigenic profile, effectively covers a majority of the circulating clades, and reaches a much higher coverage score (Fig. 2f).

### Correlation between predicted coverage score and clinical endpoints

As Fig 3a demonstrates, the coverage score has the highest correlation to vaccine effectiveness, compared to alternative metrics related to viral dominance or antigenicity. Furthermore, ranking vaccines by their coverage score allows us to distinguish between those with higher (*≥* 40) and lower (*<* 40) effectiveness (Fig. 3b). This result indicates that the coverage score holds an advantage in vaccine selection from its consideration of both collective antigenicity and the prospective dominance of viruses.

The other baselines depicted in Fig 3a illustrate that modeling viral fitness alone is insufficient for predicting vaccine effectiveness and it is critical to assess the antigenicity with the full landscape of circulating viruses. First, antigenic properties have a substantial impact on the assessment of vaccine effectiveness. Without accounting for antigenicity, the dominance of strains included in vaccines in the last season (“Dominance (last season)” in Fig. 3a) is poorly correlated with vaccine effectiveness. The antigenicity of the recommended vaccine with the most dominant virus (in the last season or predicted by EVEscape [17]) correlates better with vaccine effectiveness. Second, it is important to assess a wider viral landscape when estimating vaccine effectiveness. The strongest baseline (“Average antigenicity”) is the average negative HI value over a set of expert-selected viruses from WHO. Finally, our coverage score not only considers a collection of viruses but also incorporates their prospective dominance, demonstrating the highest correlation with vaccine effectiveness.

Beyond effectiveness, we further analyze the association between predicted coverage score and the reduction of disease burden to showcase the potential impact of our approach. The CDC estimates the reduction of disease burden via four measurements [28, 29]: the number of influenza illnesses (Fig. 3c), medical visits (Fig. 3d), hospitalizations (Fig. 3e), and deaths (Fig. 3f) averted by vaccination^3^. As shown in

Fig. 3c-e, our predicted coverage scores show positive alignment with the number of averted symptomatic illnesses, medical visits, and hospitalization (Pearson correlations of 0.6743, 0.6952, and 0.6410 respectively). Even though the coverage scores show a weaker correlation with the reduction in deaths (*R* = 0.5175), they still maintain a positive correlation. We also observed positive correlations between the estimated reduction of disease burden [28, 29] with the ground-truth coverage scores (Fig. S4) and vaccine effectiveness (Fig. S5).

### Ablation study: evaluating the superiority of the dominance and antigenicity predictor

To understand the main drivers of performance in VaxSeer, we analyze the dominance and antigenicity predictors individually. Specifically, we compared our dominance predictor against several dominance prediction baselines on the same evaluation data by fixing the antigenicity predictor.

A key aspect of VaxSeer is its ability to capture changes in dominance over time. To quantify the impact of modeling antigenic drift, we first compare our dominance predictor against a baseline that defines dominance based on the empirical frequencies calculated from the previous season (**Last**). This baseline assumes that the distribution of variants does not change between seasons. We further conduct a comparison with three machine learning models, **LM** [30], **CSCS** [15] and **EVEscape** [17], which have shown potential in predicting virus fitness and escapability based on their protein sequences [15–17]. LM uses the same architecture as our dominance predictor but predicts a static dominance without incorporating time-related information. CSCS [15] adopts the mutation probability and functional dissimilarity calculated from a masked protein language model to estimate protein fitness and escapability from antibodies. EVEscape [17] scores for individual mutations are found by combining three sources of information: a deep generative model for fitness prediction, structural information about the HA protein to estimate antibody binding potential, and chemical distances in charge and hydrophobicity between mutated and wild-type residues. In contrast to these approaches, which predict a static fitness for variants, VaxSeer adopts a dynamic perspective, learning the reproduction rate [31–33] of various virus strains and predicting their dominance accordingly. Further details on these baseline models can be found in Methods.

Another important feature of our dominance predictor is to model the entire HA protein sequence rather than single amino acid mutations [18]. Similarly to the auto-regressive natural language model GPT-2 [34], our model can indeed capture the correlation between multiple mutations (epistasis) during evolution [35]. To emphasize the importance of modeling epistasis, we compare our model with an **AA sub** model, which models the frequency evolution of single amino acid mutations.

Our results show that our proposed dominance predictor achieves the highest correlation with the ground truth coverage score, in conjunction with our antigenicity predictor (Fig. 4a and Fig. 4b). In particular, these results highlight the importance of considering the antigenic drift between seasons. While the baseline protein language models are trained using the same sequences as our model, they exhibit lower performance in dominance prediction because they are not designed to explicitly model the evolving dynamics of the dominance. In contrast, our model’s ability to capture changes in the prevalence of variants is beneficial for selecting better vaccine strains. Notably, our model also performs significantly better when incorporating interactions between mutations within HA protein sequences.

**Fig. 4:**
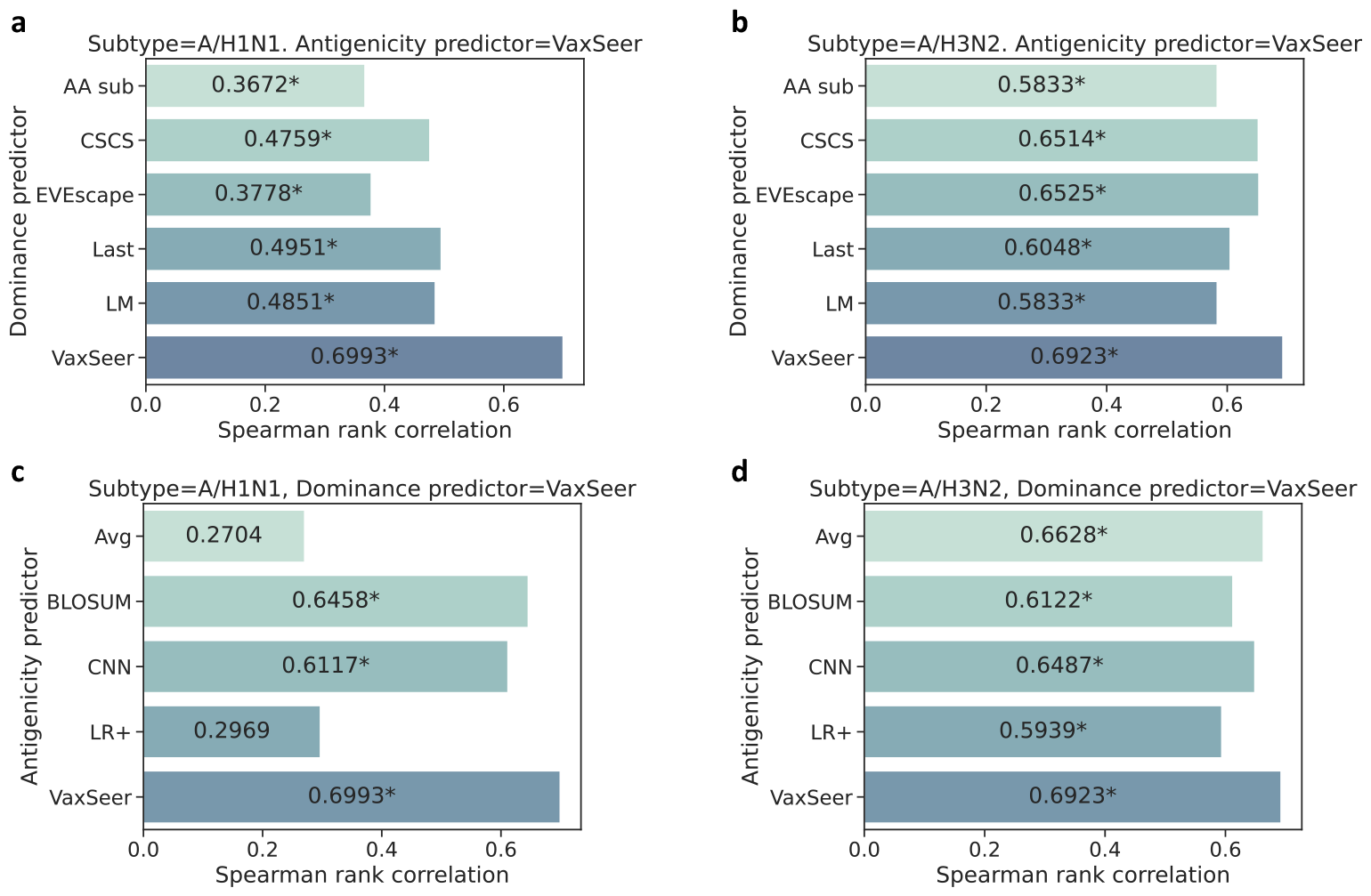
Performance of coverage score prediction. (**a, b**) Spearman rank correlation between the ground-truth coverage scores and predicted coverage scores based on different dominance predictors for A/H1N1 and A/H3N2. Given our antigenicity predictor, our dominance model achieves the best performance. (**c, d**) Spearman rank correlation between the ground-truth coverage scores and predicted coverage scores based on different antigenicity predictors for A/H1N1 and A/H3N2. Given our dominance predictor, our antigenicity predictor achieves the best performance. The star mark (*) indicates *P <* 0.01.

We next evaluate the quality of our antigenicity predictor, by fixing the dominance predictor and evaluating our antigenicity predictor against several baselines. In this setting, models take as input the aligned HA sequences from vaccines and virus strains and predict their antigenicity, i.e., the negative HI value.

Given the limited capacity of the HI test, we first measure the importance of inferring absent HI values for untested vaccine-virus combinations for scoring vaccines. We compare with a baseline, annotated as **Avg**, using experimental data collected eight months prior to the forecasted influenza season. For vaccine-virus pairs that are untested, the absent HI value is filled by the average of all experimental HI values. A sequence similarity-based imputation method is also compared, annotated as **BLO-SUM**. This method imputes the HI values for untested vaccine-virus pairs, by the results from the most similar vaccine-virus pairs that have been tested. The protein similarity is determined by the BLOSUM62 [36] matrix.

We next compared our antigenicity predictor with previous research applying traditional machine learning methods [21, 22] and other deep learning models [23, 24] to this problem. One representative baseline is a linear regression model, denoted as **LR+**[22]. It is constructed using features such as amino acid substitutions between vaccine and virus sequences, the avidity of the virus, and the potency of antiserum. The other representative baseline utilizes convolutional neural networks [37] (**CNN**) built on aligned amino acid HA sequences of viruses and vaccines. In contrast, our antigenicity predictor leveraged the MSA-Transformer [38] architecture, which is particularly well suited to the task of predicting the similarity between two input sequences.

As illustrated in Fig. 4c and Fig. 4d, our antigenicity predictor outperformed all other approaches when combined with our dominance predictor to calculate the coverage score. Specifically, our model outperformed both the LR+ and CNN models by better capturing the correlations between amino acids within and between vaccines and virus strains. In addition, we noted that the performance decreases for both imputation baselines. This indicates that the experimental results could not cover a sufficient number of circulating viruses for accurate coverage score estimation. Finally, we also found that our antigenicity predictor model was able to reconstruct the escaping mutations identified in the mutational antigenic profiling assay [20] concerning specific antibodies, even without integrating the knowledge of antibodies into our training procedure. These results are further discussed in the Supplementary Results.

## Discussion

Vaccines are an important defense against infectious diseases. However, in the presence of continual evolution, it is challenging to forecast the future landscape of viral strains and to assess the antigenicity of candidate vaccines at scale. In this study, we developed VaxSeer that selects vaccine strains based on the predicted coverage score, a prospective and holistic estimation of a vaccine’s future effectiveness. We demonstrated that VaxSeer accurately predicts the coverage score, and that vaccines selected based on these predictions have higher (retrospective) ground-truth coverage scores than the WHO’s recommendations. Furthermore, we found a statistically significant correlation between the coverage score and both vaccine effectiveness and reduced disease burden. Finally, we showed that the individual components of VaxSeer (the antigenicity predictor and the dominance predictor) outperform existing methods in their respective areas.

Though we retrospectively validated VaxSeer on an influenza case study, this approach can be applied to other neutralization assays, vaccine types, or antigens, provided that there exists data relating the genotypes of viruses and vaccines to their neutralization behavior. For instance, one could define antigenicity in terms of other influenza-related assays like HINT [39], or experiments related to other types of vaccines, e.g. mRNA vaccines. Beyond influenza, one could also apply VaxSeer to vaccine selection for other pandemic viruses like SARS-CoV-2. In this case, the dominance prediction model might be trained on Spike protein sequences with annotated collection times, and the antigenicity predictor could be trained on neutralization assays like ELISA.

This study is complementary to existing practices for vaccine selection [40], which include a variety of experimental and computational techniques for quantifying antigenicity and dominance. For example, improving wet-lab methods [8, 39] for validating vaccine candidates not only enables better assessment of those candidates but also yields a more robust and accurate antigenicity predictor. Likewise, our dominance predictor builds off existing surveillance data to forecast future trends. Compared to these experimental efforts in isolation, however, VaxSeer can efficiently explore a much larger vaccine space. For example, VaxSeer can score 100 vaccines against *∼*500 viruses within thirty minutes on a single A100 GPU. Furthermore, VaxSeer only relies on sequence data, allowing us to to compute the coverage scores for *any* vaccines or virus strains, including those that have not yet been isolated, or do not yet exist in nature. Thus, beyond its screening capabilities, VaxSeer can also be used as a tool for functional optimization and *de-novo* vaccine design.

Multiple directions can further improve this work. First, the current implementation of VaxSeer only considers the influenza virus’s HA protein, while studies have shown that the neuraminidase (NA) protein may also influence viral fitness and vaccine antigenicity [41–43]. Thus, we expect that modeling larger portions of a viral genome will provide a more complete picture for vaccine selection. Second, we only evaluated the performance of VaxSeer over vaccines with sufficient experimental antigenicity data. There exist vaccine strains with higher predicted coverage scores, but we lacked the experimental resources to validate their true antigenicity. Finally, our current approach only computes the coverage score over observed viral sequences, without considering novel sequences that may appear. In fact, during a given flu season, over 40% of HA sequences were unseen the previous year (Fig. S6). To account for these emergent sequences, we could leverage our dominance predictor as a generative model and sample viral sequences given by the predicted future distribution. This would enable us to compute the coverage score over both current and potential viral strains. Unfortunately, as stated above, today we do not have sufficient experimental data to validate the prospective effectiveness of these generative approaches.

This approach has limitations that should be acknowledged. While many factors contribute to the ultimate viability of a vaccine, this study focuses solely on antigenicity as a measure of vaccine effectiveness. Here, we do not consider other aspects of vaccine design such as host genetic background, vaccine platform, egg-adaptations in the production process, and a subject’s history of previous vaccines [44, 45]. Furthermore, our models require sufficient training data to perform well, so these architectures may encounter limited success on emergent or rare viruses. However, the overall concept of computationally forecasting antigenicity and dominance may be viable with other architectures. Finally, we only consider viral sequences isolated from human hosts, without modeling cross-species interactions. The potential impact of zoonotic viruses on vaccine effectiveness remains unexplored.

To facilitate further algorithmic development and experimental validation, our models will be made publicly available.

## Methods

### Data availability

The hemagglutination sequences and their metadata including collection time and strain name are from the GISAID (https://gisaid.org/). The hemag-glutination inhibition titer results are collected from the reports of the Worldwide Influenza Centre lab to WHO (https://www.crick.ac.uk/research/platforms-and-facilities/worldwide-influenza-centre/annual-and-interim-reports).

The human influenza vaccine composition is found from https://gisaid.org/resources/human-influenza-vaccine-composition/. The vaccine effectiveness in U.S. is from the study of CDC https://www.cdc.gov/flu/vaccines-work/effectiveness-studies.htm. The estimated influenza illnesses, medical visits, and hospitalizations averted by vaccination are found from https://www.cdc.gov/flu/vaccines-work/burden-averted.htm.

### Code availability

All models and code used for data processing, training, and evaluating VaxSeer are publicly available at https://github.com/wxsh1213/vaxseer.

### Antigenicity predictor

The hemagglutination inhibition titer (HI test) data were extracted from the published reports^4^ that have been prepared for the WHO annual consultation on the composition of influenza vaccines from 2003 to February 2023. The HI test data comprises the names of viruses and vaccine strains, along with the dilution of antibodies leading to hemagglutination inhibition. We retrieved the HA sequences with respect to strain names from GISAID. The HA protein sequences of the vaccine and testing circulating virus were aligned by the MMSeqs2 [46]. Strains for which HA sequences could not be found were excluded from the dataset. If multiple HA sequences corresponded to one strain name, we enumerated all the possible HA sequences. If a pair of vaccine-virus sequences had multiple HI test values, their geometric mean is used. We obtained the HI titer results for 70,631 distinct vaccine-virus HA protein pairs for A/H1N1 and 63,299 distinct pairs for A/H3N2.

Following previous work [22], we quantify antigenic similarity *h*(*v, x*) by the negative of relative reduction of dilution:

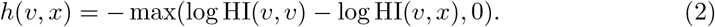

where HI(*v, x*) is the dilution of antibodies from vaccine *v* against testing strain *x*. A higher relative reduction of dilution indicates lower antibody efficacy and lower antigenic similarity between the vaccine and testing strains.

In our ablation studies, we considered both experimentally-derived and simple machine learning baselines. For the vaccine-virus pairs with available experimental HI results eight months before the winter season starts, we use their experimental results. For the vaccine-virus pairs without available experimental HI results, we use the average HI values of the whole dataset as their HI values (**Avg**). For the vaccine-virus pairs without available experimental HI results, we search for the most similar vaccine-virus pairs that have tested the HI values in the dataset and use the average of them as the estimated HI results (**BLOSUM**). The similarity is calculated from the BLOSUM62 matrix, and the similarity between two virus-vaccine pairs is the summation of the similarity between virus sequences and the similarity between vaccine sequences.

In addition, we also considered two machine learning baselines. (**LR+**) is a linear regression model whose input features are the amino acid substitution of virus and vaccine protein sequences, as well as their amino acids in each position. Finally, **CNN** is a Convolution Neural Network that consists of two 1D convolution layers with 64 channels and kernel sizes equal to 15 intervening by two one-dimensional max pooling layers. The input is the concatenation of two one-hot representations of HA protein sequences from vaccine and virus. We use the hyper-parameters suggested in [24].

### Dominance predictor

Given an amino acid sequence *x ∈ V* ^*L*^ with length *L* (where *V* is the set of all possible amino acids, typically |*V* | = 20), its probability of occurrence in season *t* is defined as

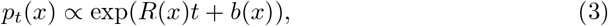

in which *R*(*x*) ∈ ℝ defines the reproduction rate (fitness) of sequence *x*, while *b*(*x*) ∈ ℝ describes when strain *x* emerges. To normalize the probability over the combinatorial space of potential sequences, we factorize the probability *p*_*t*_(*x*) autoregressively and model the conditional probability with a protein language model

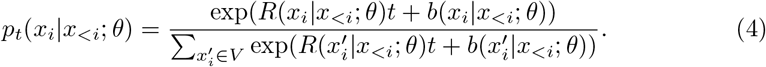

Both *R*(*x*_*i*_|*x*_*<i*_; *θ*) and *b*(*x*_*i*_|*x*_*<i*_; *θ*) are parameterized by Transformers-based deep neural networks. During the training process, we sample pairs of protein sequences and their collection times. The parameters *θ* are optimized based on the maximum likelihood estimation objective:

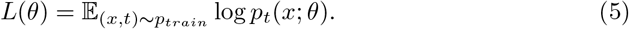

The training corpora of the dominance prediction model were obtained from the GISAID. We downloaded 394,090 HA sequences submitted before 2023-03-02. We retained only HA amino acid sequences from human hosts and with full length (with a minimum length of 553 amino acids). Starting from 2003-10, we discretized every six months into one season. Two subtypes, A/H1N1 and A/H3N2 are considered separately. For each subtype, the seasons with less than 100 HA samples are discarded. After the pre-processing, 28,546 non-repeated HA sequences are obtained for A/H3N2, and 23,736 non-repeated HA sequences are obtained for A/H1N1. The sequences were further split into training and testing sets based on collection time. Specifically, to predict the dominance of sequences in the winter season of a particular year (e.g. test set 2018-10 to 2019-04), we train on sequence data collected before the February of that year (e.g. before 2018-02).

We compared the dominance predictor to several baselines that estimate future strain dominance. Beyond those described in the main text, we provide further details on two methods. **AA Sub** models the dominance of sequences based on the dynamic likelihood of amino-acid substitutions. The probability of amino acid sequences *x* occurring in season *t* is the product of the probabilities of amino acid at each position, considered independently.

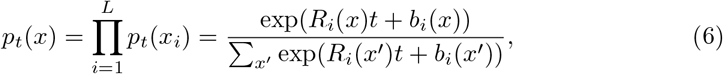

in which *R*_*i*_(*x*) and *b*_*i*_(*x*) are learnable scalar parameters describing the reproduction rate and emerging time for amino acid *x* at site *i*.

**LM** models the dominance of sequences in a static manner. Similarly to Eq. 4, the likelihood of amino acid sequence *x* is factorized autoregressively without using the collection time information:

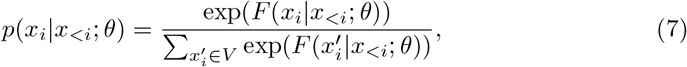

in which *F* (*x*_*i*_|*x*_*<i*_; *θ*) is the logits output by a GTP-2 [34].

## Supplementary materials

### Interpreting antigenicity predictor with validated escaping mutations

**Fig. 1:**
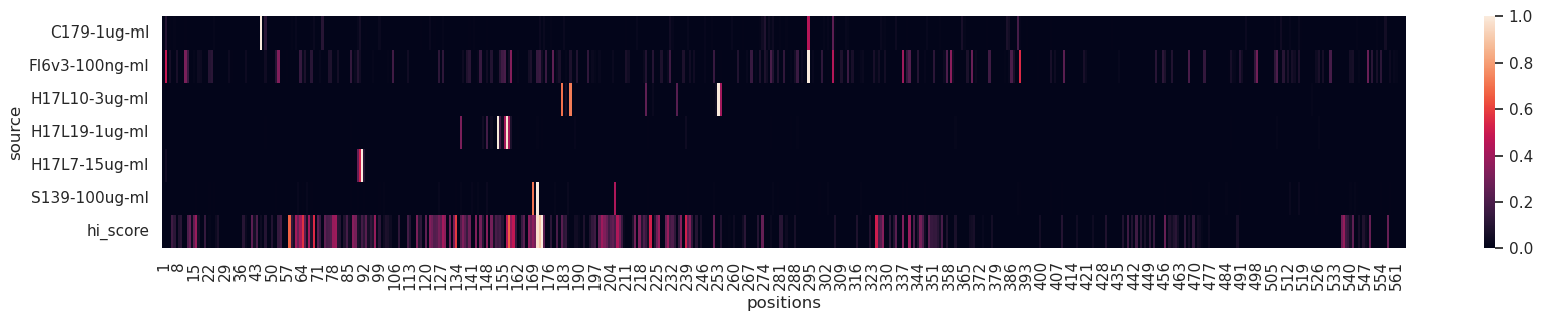
Escapable effects of single amino acid mutations given by antigenicity predictor and mutational antigenic profiling assays relative to different antibodies. The *x*-axis is the position in reference virus strain A/WSN/1933 of A/H1N1. The mutational antigenic profiling results are from [20].

Our antigenicity predictor could identify the escaping mutations observed in other neutralization assays with respect to specific antibodies, without incorporating any of this information into our training process. The mutational antigenic profiling assay [20] investigates the effect of single amino acid mutations on escaping broad and narrow antibodies targeting HA. We predict the HI value between the same wild-type vaccine strain A/WSN/1933 used in their experiment and the same virus strains with single amino acid mutations to the wild-type strain. The quantified escape effect given by the mutational antigenic profiling assays and our model is shown in Fig. 1. Our model predicts that the mutations on site 171 allow for escape from the antibodies produced by A/WSN/1933. This is consistent with the discovery in mutational antigenic profiling results that mutations in site 171 allow for escape from the broadly neutralizing antibody S139 [20]. Our model also predicts that mutations on site 158 escape from the vaccine, which is consistent with the escaping mutations from antibody H17L19 given by mutational antigenic profiling assays [20]. Finally, our model assigned high HI values to some additional sites, but mutations at these points have not been observed to escape from the antibodies screened thus far. Additional experiments are required to elucidate these sites, which might escape antibodies that have not yet been identified, or act through unknown immune escape mechanisms.

**Fig. 2:**
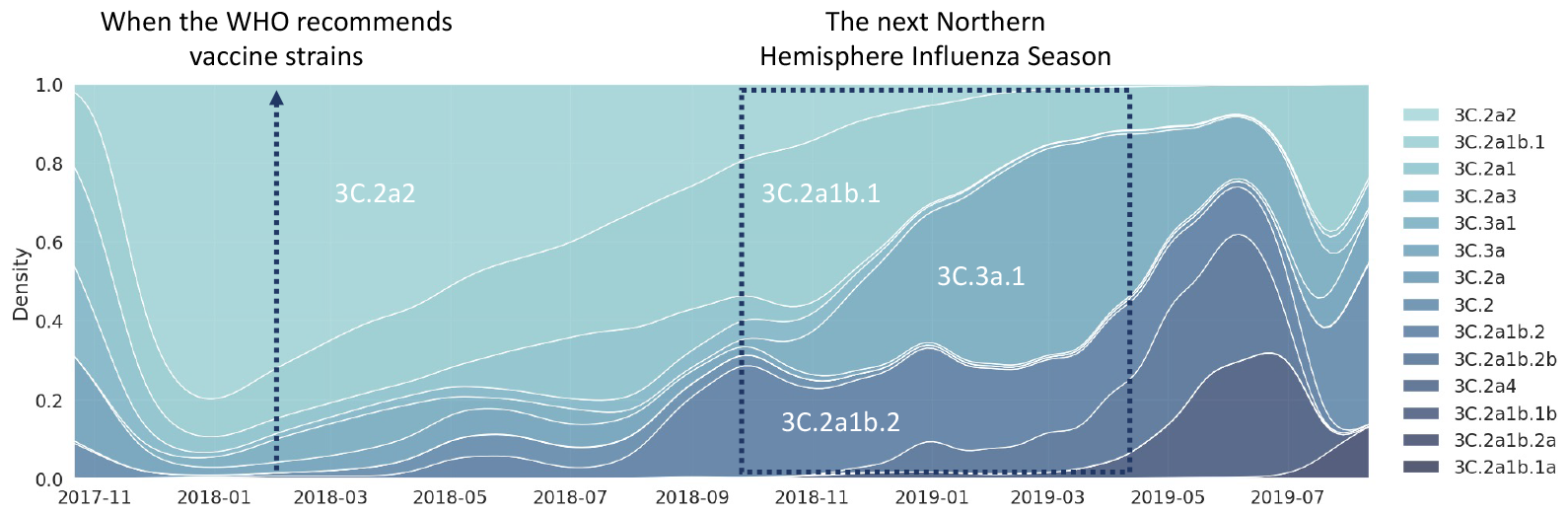
Antigenic drift of A/H3N2 occurs between the period of vaccine selection and the following Northern Hemisphere Influenza Season. Prior to the vaccine strain selection, the prevailing clade was 3C.2a2. However, in the subsequent influenza season, the dominant clades shift to 3C.2a1b.1, 3C.3a.1, and 3C.2a1b.2. The density of each clade is calculated by aggregating the occurrence of HA sequences belonging to that clade. HA sequences are obtained from GISAID, and their clades are annotated by Nextclade [47].

**Fig. 3:**
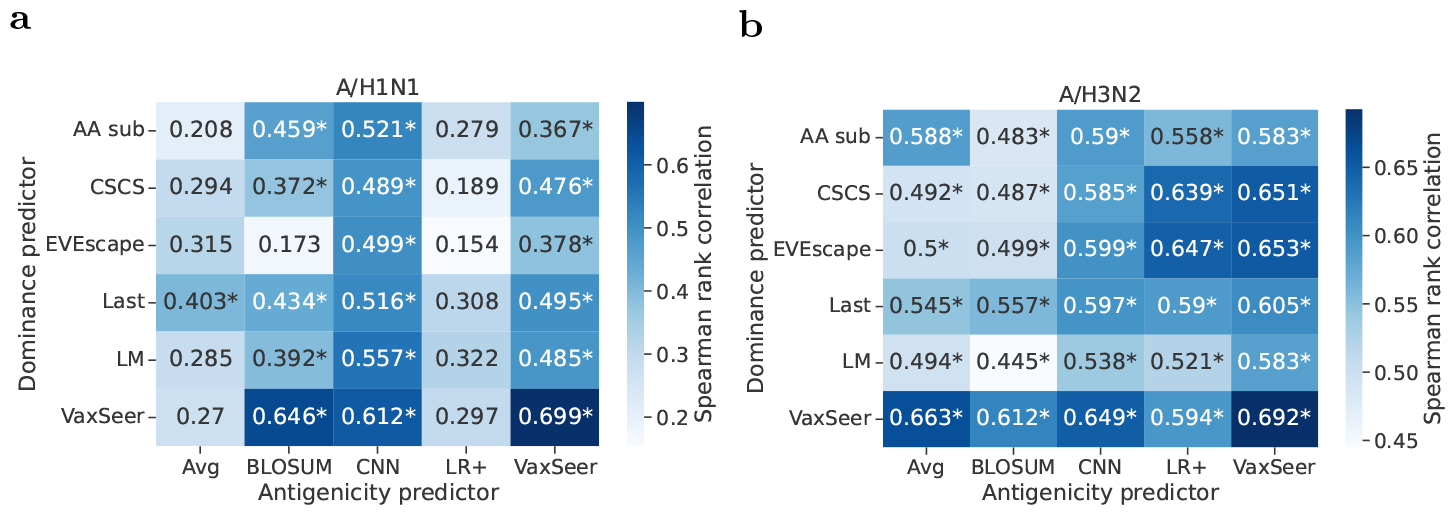
Spearman rank correlation *ρ* between the predicted coverage scores and ground-truth scores (sequence-level) for A/H3N2 and A/H1N1, from 2012 to 2021 seasons. Vaccines with HI data covering more than 40% circulating sequences are compared. The star mark (*) indicates *P <* 0.01.

**Fig. 4:**
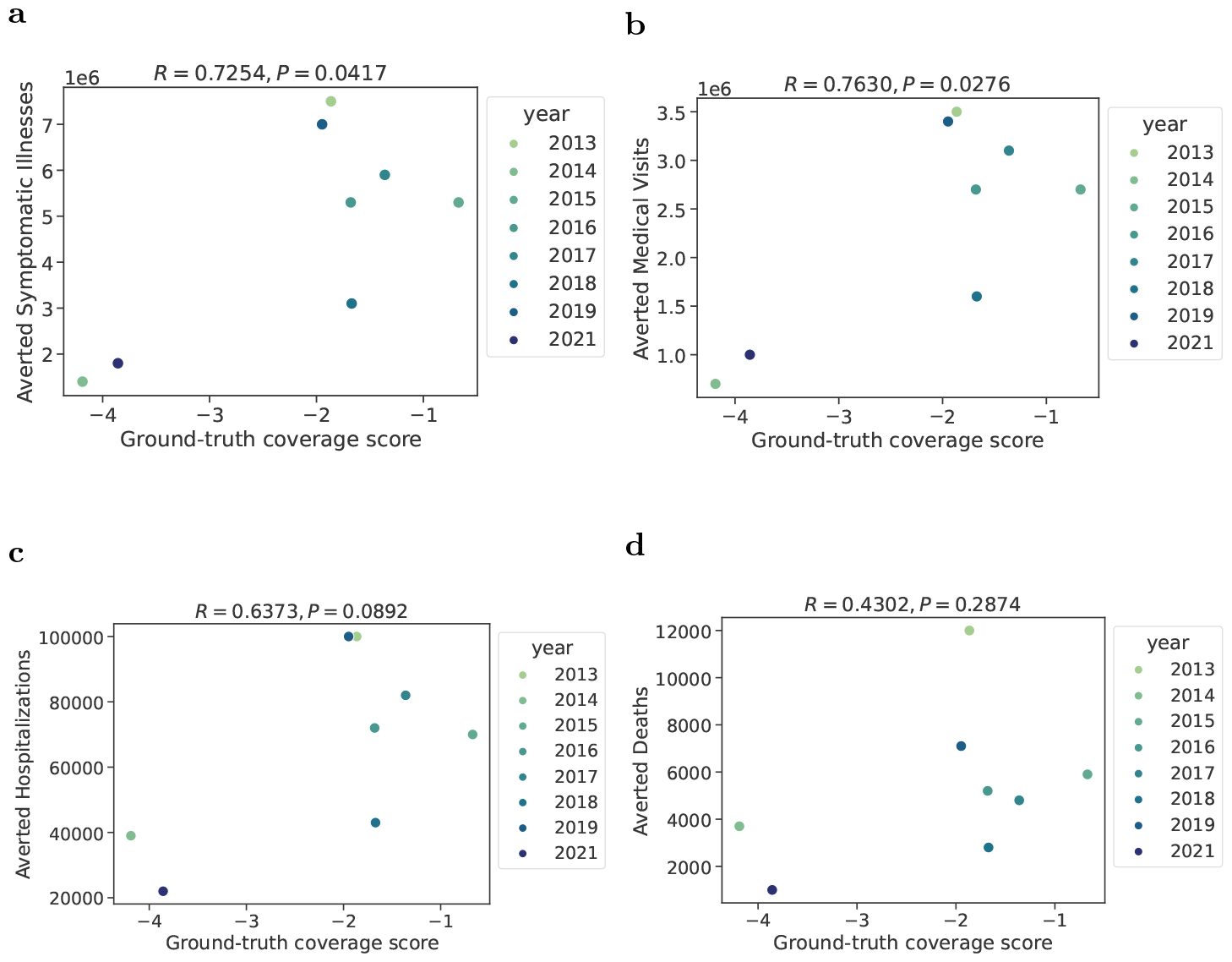
The ground-truth coverage score is correlated with disease burden prevention. Ground-truth coverage scores and the estimated number of influenza illnesses (**a**), medical visits (**b**), hospitalizations (**c**), and deaths (**d**) averted by vaccination. Pearson correlations (*R*) and corresponding P-values (*P*) are illustrated in the text.

**Fig. 5:**
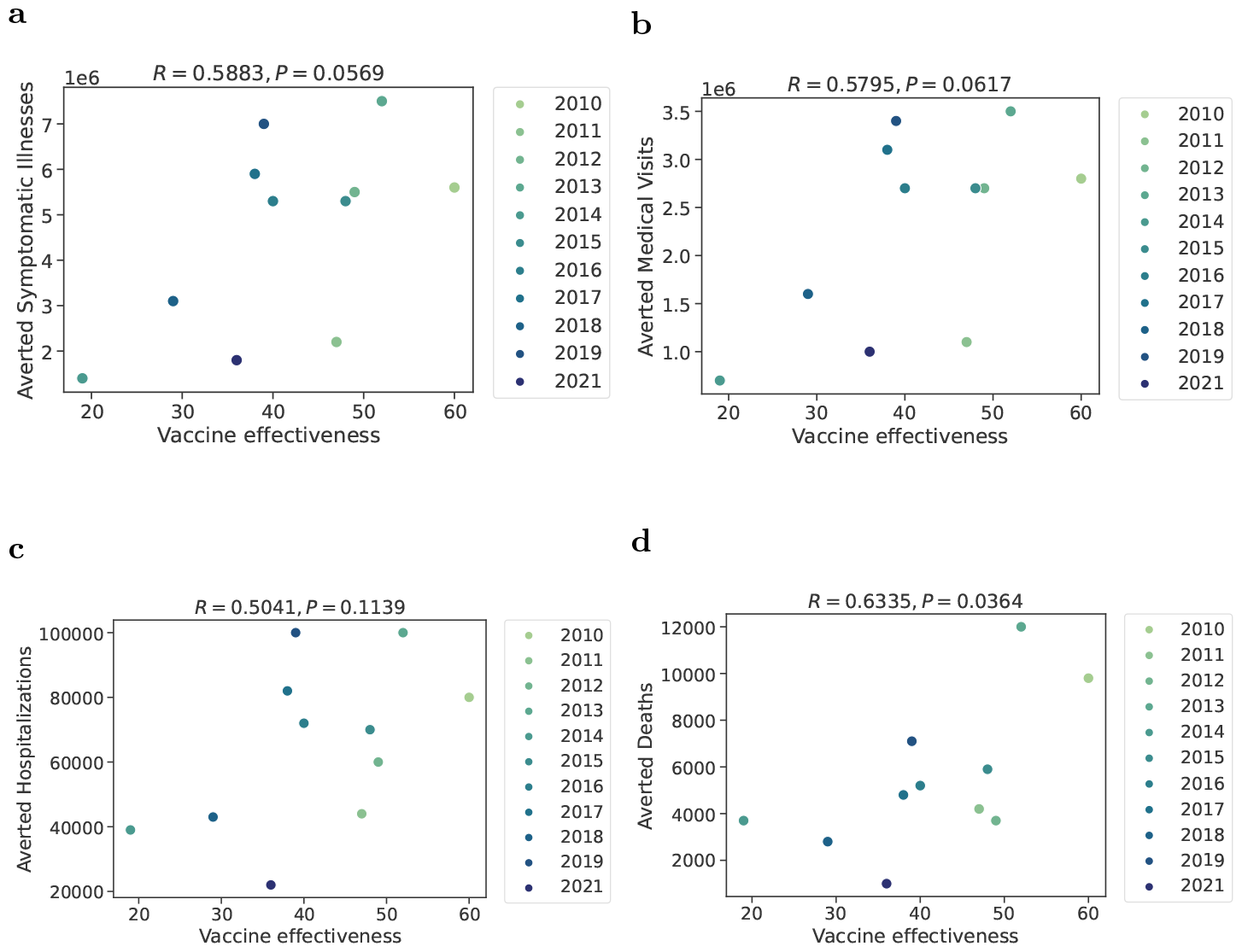
The vaccine effectiveness is correlated with disease burden prevention. The vaccine effectiveness and the estimated number of influenza illnesses (**a**), medical visits (**b**), hospitalizations (**c**), and deaths (**d**) averted by vaccination. Pearson correlations (*R*) and corresponding P-values (*P*) are illustrated in the text.

**Fig. 6:**
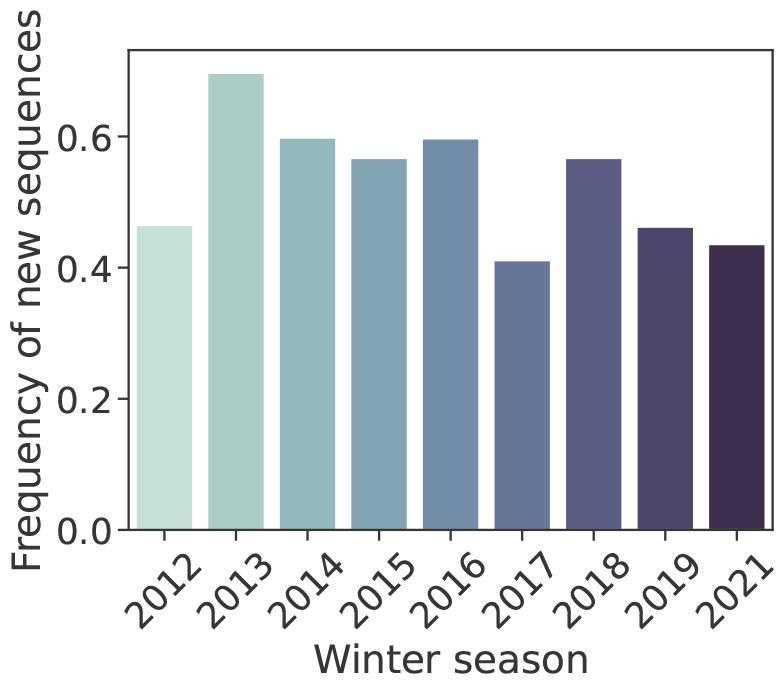
The frequency of new HA sequences occurring in each winter season for A/H3N2. The frequency is calculated from the sequence data with collection time obtained from GISAID.

## Footnotes

1 Influenza vaccine effectiveness (VE) is defined as the reduction in the odds of influenza infection among vaccinated individuals compared to those who are not vaccinated. The VE used in this paper is estimated by test-negative design [1] with study subjects seeking care for an acute respiratory illness (ARI).

2 https://www.crick.ac.uk/research/platforms-and-facilities/worldwide-influenza-centre/annual-and-interim-reports

3 https://www.cdc.gov/flu/vaccines-work/burden-averted.htm

4 https://www.crick.ac.uk/research/platforms-and-facilities/worldwide-influenza-centre/annual-and-interim-reports

## References

[1] Jackson, M.L., Nelson, J.C.: The test-negative design for estimating influenza vaccine effectiveness. Vaccine 31(17), 2165–2168 (2013)

[2] Dawood, F.S., Chung, J.R., Kim, S.S., Zimmerman, R.K., Nowalk, M.P., Jackson, M.L., Jackson, L.A., Monto, A.S., Martin, E.T., Belongia, E.A., et al.: Interim estimates of 2019–20 seasonal influenza vaccine effectiveness—united states, february 2020. Morbidity and Mortality Weekly Report 69(7), 177 (2020)

[3] Zimmerman, R.K., Nowalk, M.P., Chung, J., Jackson, M.L., Jackson, L.A., Petrie, J.G., Monto, A.S., McLean, H.Q., Belongia, E.A., Gaglani, M., et al.: 2014– 2015 influenza vaccine effectiveness in the united states by vaccine type. Clinical Infectious Diseases, 635 (2016)

[4] Rolfes, M.A., Flannery, B., Chung, J.R., O’Halloran, A., Garg, S., Belongia, E.A., Gaglani, M., Zimmerman, R.K., Jackson, M.L., Monto, A.S., et al.: Effects of influenza vaccination in the united states during the 2017–2018 influenza season. Clinical Infectious Diseases 69(11), 1845–1853 (2019)

[5] Flannery, B., Kondor, R.J.G., Chung, J.R., Gaglani, M., Reis, M., Zimmerman, R.K., Nowalk, M.P., Jackson, M.L., Jackson, L.A., Monto, A.S., et al.: Spread of antigenically drifted influenza a (h3n2) viruses and vaccine effectiveness in the united states during the 2018–2019 season. The Journal of infectious diseases 221(1), 8–15 (2020)

[6] Tenforde, M.W., Kondor, R.J.G., Chung, J.R., Zimmerman, R.K., Nowalk, M.P., Jackson, M.L., Jackson, L.A., Monto, A.S., Martin, E.T., Belongia, E.A., et al.: Effect of antigenic drift on influenza vaccine effectiveness in the united states—2019–2020. Clinical Infectious Diseases 73(11), 4244–4250 (2021)

[7] Price, A.M., Flannery, B., Talbot, H.K., Grijalva, C.G., Wernli, K.J., Phillips, C.H., Monto, A.S., Martin, E.T., Belongia, E.A., McLean, H.Q., et al.: Influenza vaccine effectiveness against influenza a (h3n2)-related illness in the united states during the 2021–2022 influenza season. Clinical Infectious Diseases 76(8), 1358–1363 (2023)

[8] Hirst, G.K.: Studies of antigenic differences among strains of influenza a by means of red cell agglutination. The Journal of experimental medicine 78(5), 407–423 (1943)

[9] Barr, I.G., Russell, C., Besselaar, T.G., Cox, N.J., Daniels, R.S., Donis, R., Engelhardt, O.G., Grohmann, G., Itamura, S., Kelso, A., et al.: Who recommendations for the viruses used in the 2013–2014 northern hemisphere influenza vaccine: Epidemiology, antigenic and genetic characteristics of influenza a (h1n1) pdm09, a (h3n2) and b influenza viruses collected from october 2012 to january 2013. Vaccine 32(37), 4713–4725 (2014)

[10] Gerdil, C.: The annual production cycle for influenza vaccine. Vaccine 21(16), 1776–1779 (2003)

[11] Smith, D.J., Lapedes, A.S., De Jong, J.C., Bestebroer, T.M., Rimmelzwaan, G.F., Osterhaus, A.D., Fouchier, R.A.: Mapping the antigenic and genetic evolution of influenza virus. science 305(5682), 371–376 (2004)

[12] Gouma, S., Weirick, M., Hensley, S.E.: Antigenic assessment of the h3n2 component of the 2019-2020 northern hemisphere influenza vaccine. Nature Communications 11(1), 2445 (2020)

[13] Flannery, B., Kondor, R.J.G., Chung, J.R., Gaglani, M., Reis, M., Zimmerman, R.K., Nowalk, M.P., Jackson, M.L., Jackson, L.A., Monto, A.S., et al.: Spread of antigenically drifted influenza a (h3n2) viruses and vaccine effectiveness in the united states during the 2018–2019 season. The Journal of infectious diseases 221(1), 8–15 (2020)

[14] Luksza, M., Lässig, M.: A predictive fitness model for influenza. Nature 507(7490), 57–61 (2014)

[15] Hie, B., Zhong, E.D., Berger, B., Bryson, B.: Learning the language of viral evolution and escape. Science 371(6526), 284–288 (2021)

[16] Frazer, J., Notin, P., Dias, M., Gomez, A., Min, J.K., Brock, K., Gal, Y., Marks, D.S.: Disease variant prediction with deep generative models of evolutionary data. Nature 599(7883), 91–95 (2021)

[17] Thadani, N.N., Gurev, S., Notin, P., Youssef, N., Rollins, N.J., Sander, C., Gal, Y., Marks, D.S.: Learning from pre-pandemic data to forecast viral escape. Nature (2023)

[18] Obermeyer, F., Jankowiak, M., Barkas, N., Schaffner, S.F., Pyle, J.D., Yurkovetskiy, L., Bosso, M., Park, D.J., Babadi, M., MacInnis, B.L., et al.: Analysis of 6.4 million sars-cov-2 genomes identifies mutations associated with fitness. Science 376(6599), 1327–1332 (2022)

[19] Han, W., Chen, N., Xu, X., Sahil, A., Zhou, J., Li, Z., Zhong, H., Gao, E., Zhang, R., Wang, Y., et al.: Predicting the antigenic evolution of sars-cov-2 with deep learning. Nature Communications 14(1), 3478 (2023)

[20] Doud, M.B., Lee, J.M., Bloom, J.D.: How single mutations affect viral escape from broad and narrow antibodies to h1 influenza hemagglutinin. Nature communications 9(1), 1386 (2018)

[21] Zeller, M.A., Gauger, P.C., Arendsee, Z.W., Souza, C.K., Vincent, A.L., Anderson, T.K.: Machine learning prediction and experimental validation of antigenic drift in h3 influenza a viruses in swine. MSphere 6(2), 10–1128 (2021)

[22] Neher, R.A., Bedford, T., Daniels, R.S., Russell, C.A., Shraiman, B.I.: Prediction, dynamics, and visualization of antigenic phenotypes of seasonal influenza viruses. Proceedings of the National Academy of Sciences 113(12), 1701–1709 (2016)

[23] Yin, R., Thwin, N.N., Zhuang, P., Lin, Z., Kwoh, C.K.: Iav-cnn: a 2d convolutional neural network model to predict antigenic variants of influenza a virus. IEEE/ACM Transactions on Computational Biology and Bioinformatics 19(6), 3497–3506 (2021)

[24] Xia, Y.-L., Li, W., Li, Y., Ji, X.-L., Fu, Y.-X., Liu, S.-Q., et al.: A deep learning approach for predicting antigenic variation of influenza a h3n2. Computational and mathematical methods in medicine 2021 (2021)

[25] Krammer, F.: The human antibody response to influenza a virus infection and vaccination. Nature Reviews Immunology 19(6), 383–397 (2019)

[26] Gamblin, S.J., Skehel, J.J.: Influenza hemagglutinin and neuraminidase membrane glycoproteins. Journal of biological chemistry 285(37), 28403–28409 (2010)

[27] Khare, S., Gurry, C., Freitas, L., Schultz, M.B., Bach, G., Diallo, A., Akite, N., Ho, J., Lee, R.T., Yeo, W., et al.: Gisaid’s role in pandemic response. China CDC weekly 3(49), 1049 (2021)

[28] Kostova, D., Reed, C., Finelli, L., Cheng, P.-Y., Gargiullo, P.M., Shay, D.K., Singleton, J.A., Meltzer, M.I., Lu, P.-j., Bresee, J.S.: Influenza illness and hospitalizations averted by influenza vaccination in the united states, 2005–2011. PloS one 8(6), 66312 (2013)

[29] Reed, C., Chaves, S.S., Daily Kirley, P., Emerson, R., Aragon, D., Hancock, E.B., Butler, L., Baumbach, J., Hollick, G., Bennett, N.M., et al.: Estimating influenza disease burden from population-based surveillance data in the united states. PloS one 10(3), 0118369 (2015)

[30] Ferruz, N., Schmidt, S., Höcker, B.: Protgpt2 is a deep unsupervised language model for protein design. Nature communications 13(1), 4348 (2022)

[31] Kermack, W.O., McKendrick, A.G.: A contribution to the mathematical theory of epidemics. Proceedings of the royal society of london. Series A, Containing papers of a mathematical and physical character 115(772), 700–721 (1927)

[32] Harko, T., Lobo, F.S., Mak, M.: Exact analytical solutions of the susceptible-infected-recovered (sir) epidemic model and of the sir model with equal death and birth rates. Applied Mathematics and Computation 236, 184–194 (2014)

[33] Beckley, R., Weatherspoon, C., Alexander, M., Chandler, M., Johnson, A., Bhatt, G.S.: Modeling epidemics with differential equations. Tennessee State University Internal Report (2013)

[34] Radford, A., Wu, J., Child, R., Luan, D., Amodei, D., Sutskever, I.: Language models are unsupervised multitask learners (2019)

[35] Gong, L.I., Suchard, M.A., Bloom, J.D.: Stability-mediated epistasis constrains the evolution of an influenza protein. Elife 2, 00631 (2013)

[36] Henikoff, S., Henikoff, J.G.: Amino acid substitution matrices from protein blocks. Proceedings of the National Academy of Sciences 89(22), 10915–10919 (1992)

[37] Kiranyaz, S., Avci, O., Abdeljaber, O., Ince, T., Gabbouj, M., Inman, D.J.: 1d convolutional neural networks and applications: A survey. Mechanical systems and signal processing 151, 107398 (2021)

[38] Rao, R.M., Liu, J., Verkuil, R., Meier, J., Canny, J., Abbeel, P., Sercu, T., Rives, A.: Msa transformer. In: International Conference on Machine Learning, pp. 8844–8856 (2021). PMLR

[39] Jorquera, P.A., Mishin, V.P., Chesnokov, A., Nguyen, H.T., Mann, B., Garten, R., Barnes, J., Hodges, E., De La Cruz, J., Xu, X., et al.: Insights into the antigenic advancement of influenza a (h3n2) viruses, 2011–2018. Scientific Reports 9(1), 2676 (2019)

[40] Engelhardt, O., Hasegawa, H., Lewis, N., Subbarao, K., Wang, D., Webby, R., Wentworth, D., Ye, Z.a.: Recommended composition of influenza virus vaccines for use in the 2023-2024 northern hemisphere influenza season (2023)

[41] Matrosovich, M.N., Matrosovich, T.Y., Gray, T., Roberts, N.A., Klenk, H.-D.: Neuraminidase is important for the initiation of influenza virus infection in human airway epithelium. Journal of virology 78(22), 12665–12667 (2004)

[42] Sylte, M.J., Suarez, D.L.: Influenza neuraminidase as a vaccine antigen. Vaccines for Pandemic Influenza, 227–241 (2009)

[43] Monto, A.S., Petrie, J.G., Cross, R.T., Johnson, E., Liu, M., Zhong, W., Levine, M., Katz, J.M., Ohmit, S.E.: Antibody to influenza virus neuraminidase: an independent correlate of protection. The Journal of infectious diseases 212(8), 1191–1199 (2015)

[44] Han, A.X., Jong, S.P., Russell, C.A.: Co-evolution of immunity and seasonal influenza viruses. Nature Reviews Microbiology, 1–13 (2023)

[45] Lewnard, J.A., Cobey, S.: Immune history and influenza vaccine effectiveness. Vaccines 6(2), 28 (2018)

[46] Steinegger, M., Söding, J.: Mmseqs2 enables sensitive protein sequence searching for the analysis of massive data sets. Nature biotechnology 35(11), 1026–1028 (2017)

[47] Aksamentov, I., Roemer, C., Hodcroft, E.B., Neher, R.A.: Nextclade: clade assignment, mutation calling and quality control for viral genomes. Journal of open source software 6(67), 3773 (2021)

